# CoLa-VAE: A Cell–Cell Communication-Aware Variational Autoencoder for Representation Learning and Expression Denoising

**DOI:** 10.64898/2026.03.28.715052

**Authors:** Yeqing Chen, Cong Qi, Hanzhang Fang, Feiyang Luan, Zhirong Zhang, Shivvrat Arya, Zhi Wei

## Abstract

Single-cell RNA sequencing provides a powerful view of cellular heterogeneity, but its sparsity and dropout noise remain major obstacles for recovering biologically meaningful gene expression programs and for downstream analyses that depend on reliable expression measurements. Ligand–receptor-based cell–cell communication inference is such analysis, missing ligand or receptor expression can cause substantial false negatives in sparse single-cell data. Here, we present CoLa-VAE, a cell–cell communication-aware variational autoencoder that jointly learns latent representations and denoised expression profiles by incorporating ligand–receptor-derived communication topology through dynamic graph Laplacian regularization. Rather than treating denoising as a secondary output of representation learning, CoLa-VAE uses denoised expression to iteratively refine communication estimates and uses the resulting communication structure to guide both latent organization and expression reconstruction. In addition to improving latent space organization and producing robust denoised expression matrices, CoLa-VAE-denoised matrices also improved downstream biological analyses, including the detection of robust differential cell–cell communication programs, mitigation of batch-associated variation and enhanced spatial transcriptomic deconvolution when spatially constrained communication structure was incorporated. Together, these results establish CoLa-VAE as a communication-guided denoising and representation learning framework that recovers biologically meaningful expression signals from sparse single-cell and spatial transcriptomic data, enabling more sensitive and reliable downstream analysis.

## Introduction

Recent advances in single-cell RNA sequencing (scRNA-seq) have enabled high-resolution characterization of cellular heterogeneity across diverse biological systems. A central computational task in single-cell analysis is representation learning, which aims to compress high-dimensional gene expression measurements into a low-dimensional latent space that captures major sources of biological variation^1^. Variational inference has emerged as a particularly powerful framework for this purpose. In variational autoencoder (VAE)-based models, a tractable distribution is used to approximate the intractable posterior, enabling scalable probabilistic modeling of gene expression distributions. Methods such as scVI^2^ and related generative frameworks have demonstrated strong performance in modeling transcriptomic variability and providing embeddings that support downstream tasks including clustering, batch integration, and denoising.

Despite these advances, most existing representation learning frameworks model each cell primarily through its intrinsic transcriptional profile. Under this assumption, each cell is treated largely as an independent observation whose expression profile is generated from a cell-specific latent variable. While this strategy has been effective for capturing global transcriptional structure, it does not explicitly account for the fact that cells in multicellular systems are shaped not only by intrinsic gene regulatory programs but also by extrinsic signals received from neighboring cells through ligand–receptor-mediated communication^3,4^. These extrinsic interactions can substantially influence transcriptional states, particularly in plastic cell populations such as immune, stromal, and glial cells whose functional identities depend strongly on local niche signals.

Single-cell data also present another closely related challenge, the observed count matrix is an incomplete and noisy measurement of the underlying biological expression state. Dropout events and low capture efficiency lead to pervasive sparsity, especially for genes expressed at low or moderate levels^5^. This issue is particularly important for downstream analyses that rely on detecting coordinated expression of functionally related genes. Cell–cell communication (CCC) inference is a representative example^3^. Widely used tools such as CellChat^6^, CellPhoneDB^7^ and related methods^8^ estimate signaling interactions from ligand and receptor expression. However, when ligand or receptor expression is missed due to technical dropout, communication probabilities can be underestimated, leading to false-negative interactions. For this reason, many CCC tools operate at the celltype or cluster level, where expression is aggregated to stabilize inference, but this aggregation can obscure cell-state heterogeneity and sample-specific communication changes^9^.

These considerations create a practical chicken-and-egg problem. Accurate communication inference requires a reliable expression matrix^9^, but accurate recovery of biologically meaningful expression may itself benefit from knowledge of communication structure. Cells that participate in similar CCC signaling programs or are exposed to similar CCC signaling environments are often functionally related, even when their raw expression profiles are noisy or partially observed. Thus, communication topology may provide a complementary biological signal that can guide both representation learning and expression denoising.

To address this challenge, we developed CoLa-VAE, a cell–cell communication-aware variational autoencoder that jointly learns a latent representation and a denoised expression matrix. CoLa-VAE extends the standard VAE framework by incorporating ligand–receptor-derived communication topology into model training through a dynamic graph Laplacian constraint. During training, the decoder generates a denoised expression estimate, which is used to infer single-cell-level communication profiles across ligand–receptor pairs. Cells are then compared based on their global outgoing and incoming signaling behaviors, reflecting how similarly they send signals to other cells and how similarly they receive signals from their environment^10^. This communication-derived similarity is converted into a graph structure that regularizes the latent space, encouraging cells with similar signaling roles to occupy nearby positions.

A key feature of CoLa-VAE is that the communication graph is not fixed. Instead, communication inference and expression denoising are updated iteratively during training. As the model reconstructs a less sparse expression matrix, ligand–receptor interactions can be estimated more reliably; the resulting communication structure then provides a biological constraint that further improves latent organization and denoising. This dynamic feedback design allows CoLa-VAE to use cell–cell communication not only as an auxiliary annotation layer, but also as a functional prior for recovering expression programs that are obscured by technical noise.

Importantly, CoLa-VAE is modular with respect to the communication inference strategy. The framework can incorporate communication scores derived from different ligand–receptor-based formulations, including CellChat-, CellPhoneDB-, iTALK-, and CytoTalk-like scoring schemes. This plug-and-play design allows CoLa-VAE to remain agnostic to a specific CCC tool while leveraging the broader principle that communication similarity captures biologically meaningful relationships among cells.

In this study, we show that CoLa-VAE improves both latent representation quality and denoised expression recovery across multiple single-cell and spatial transcriptomic settings. In PBMC datasets, CoLa-VAE produced structured latent embeddings across different CCC modules and improved robustness under expression-dependent masking. In simulation experiments with known ground-truth expression, CoLa-VAE-denoised matrices more closely recovered the underlying expression state than scVI and dedicated imputation methods including MAGIC^11^ and ALRA^12^. Across PBMC datasets generated from different sequencing platforms, CoLa-VAE improved denoised expression consistency within matched cell types and partially mitigated batch-associated variation. We further demonstrate that CoLa-VAE-denoised matrices enhance downstream biological analyses, including the detection of robust differential cell–cell communication alterations in opioid use disorder dataset and improved celltype deconvolution in spatial transcriptomics when spatial constraints are incorporated.

Together, these results reposition CoLa-VAE as more than a communication-aware latent representation model. CoLa-VAE provides a communication-guided denoising and imputation framework that uses intercellular signaling structure to recover biologically meaningful expression signals from sparse single-cell data. By producing both interpretable latent embeddings and high-quality denoised expression matrices, CoLa-VAE offers a flexible strategy for improving downstream analyses that depend on reliable gene expression measurements, including clustering, batch comparison, differential communication analysis, and spatial deconvolution.

## Results

### CoLa-VAE integrates expression reconstruction with communication-derived cellular similarity

Single-cell transcriptional states are shaped not only by cell-intrinsic gene regulatory programs, but also by extrinsic signals mediated through intercellular communication (Fig. 1a). Based on this premise, we developed CoLa-VAE to incorporate communication-derived cellular relationships into variational representation learning. CoLa-VAE takes a raw scRNA-seq count matrix as input and uses ligand-receptor interaction information derived from CCC inference tools, including CellChat, CellPhoneDB, iTALK, and CytoTalk, to construct communication-aware constraints. The model jointly learns a latent representation and a denoised expression matrix, enabling downstream analyses such as clustering, differential expression analysis, and cell-cell communication analysis (Fig 1b). Conceptually, the framework first summarizes ligand-receptor interaction patterns across cells, transforms these patterns into a communication-derived similarity structure (Fig 1c), and then uses this structure to regularize the disentanglement latent space through a graph-based loss (Fig 1d).

**Figure 1.**
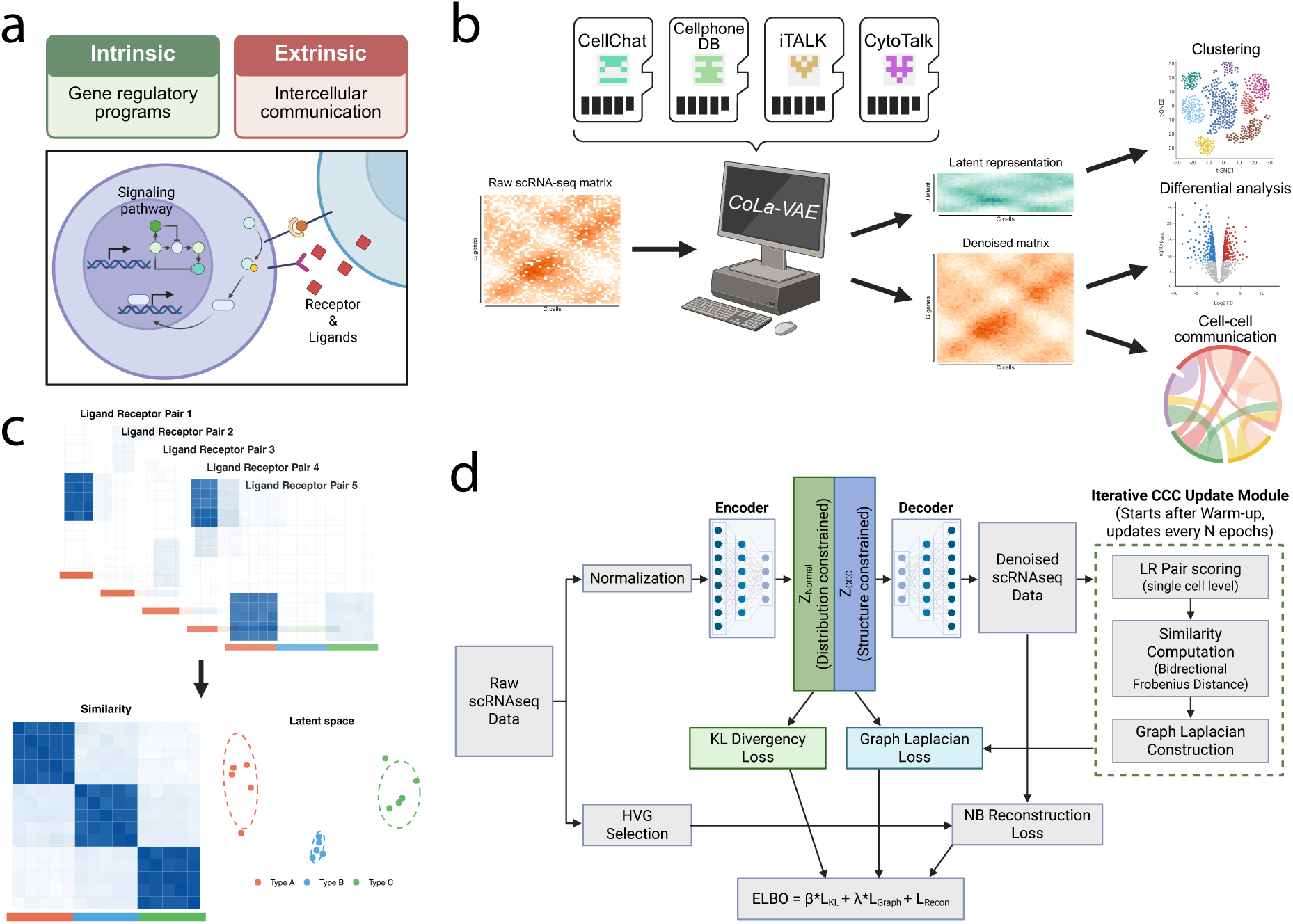
CoLa-VAE framework integrates intrinsic transcriptional programs and extrinsic cell–cell communication to learn CCC-aware representations. **(a)** Conceptual overview of cell identity as a combination of intrinsic gene regulatory programs and extrinsic intercellular communication. **(b)** Overview of the CoLa-VAE pipeline. Starting from a gene expression matrix, communication signals are estimated using a plug-in CCC module (e.g., CellChat). The model jointly learns a latent representation and a denoised expression matrix, enabling downstream analyses such as clustering, differential expression, and communication inference. **(c)** Schematic illustration of CCC-informed similarity construction. For each ligand–receptor (LR) pair, a cell–cell communication matrix is computed. These matrices are aggregated to define pairwise cell similarity based on shared CCC signaling patterns. Cells with similar CCC profiles are mapped closer in the latent space, forming structured clusters. **(d)** Model architecture of CoLa-VAE. The encoder–decoder framework reconstructs gene expression while incorporating a CCC module that computes communication-derived similarities. A graph Laplacian loss enforces that cells with similar communication profiles remain close in the latent space, alongside reconstruction and KL divergence losses. Together, these objectives yield CCC-aware latent embeddings and denoised expression profiles.

To evaluate whether communication-derived similarity captures meaningful biological structure, we first used PBMC3k as a proof-of-concept dataset, we computed cell–cell communication matrices across ligand–receptor (LR) pairs (Supplementary Fig. 1a) and quantified pairwise similarity using outgoing and incoming signaling profiles (Supplementary Fig. 1a). The resulting communication-derived distances exhibited clear biological structure. Cells belonging to the same annotated cell type showed significantly smaller outgoing, incoming, and bidirectional communication distances compared with cells from different cell types (Supplementary Fig. 1B), indicating that cells sharing similar biological identities also tend to occupy similar signaling environments.

Transformation of these communication distances into a similarity graph produced pronounced block-like structures aligned with known immune cell populations (Supplementary Fig. 1c). Cell-type-level aggregation further demonstrated that within-cell-type similarities were substantially higher than between-cell-type similarities (Supplementary Fig. 1d), supporting the hypothesis that communication topology captures biologically meaningful cellular organization. Together, these results establish that CCC-derived similarity structures provide an informative and biologically coherent constraint for latent representation learning.

### CoLa-VAE improves representation learning and denoising accuracy

We next evaluated whether incorporating communication-aware constraints improves single-cell representation learning and denoising performance. Using the PBMC3k dataset, we compared CoLa-VAE with Seurat and scVI and further tested whether the framework remained stable when different CCC inference strategies were used to define communication features. Specifically, we implemented CoLa-VAE with modified single cell version of CellChat-, CellPhoneDB-, CytoTalk- and iTALK-derived CCC formulations. Across all four formulations, CoLa-VAE generated well-organized latent embeddings that separated major immune cell populations, including B cells, CD14+ monocytes, FCGR3A+ monocytes, dendritic cells, CD4 T cells, CD8 T cells, NK cells, and platelets (Fig. 2a). The overall organization of the latent space was largely preserved across CCC modules, suggesting that CoLa-VAE is not dependent on a single CCC inference method. We next quantified the representation quality using both clustering-structure metrics and annotation-based metrics. Across silhouette index (SI), Dunn index (DI), Calinski-Harabasz index (CHI), normalized mutual information (NMI), adjusted Rand index (ARI), and macro-F1 score, CoLa-VAE models with different CCC modules generally achieved better performance than Seurat and scVI (Fig. 2b). When these metrics were aggregated into a mean normalized score, all CoLa-VAE variants ranked above the baseline methods, with CellPhoneDB-, CytoTalk-, CellChat-, and iTALK-derived implementations showing similar overall performance (Fig. 2c). These results indicate that communication-aware constraints improve latent representation while remaining robust to the choice of CCC modules.

**Figure 2.**
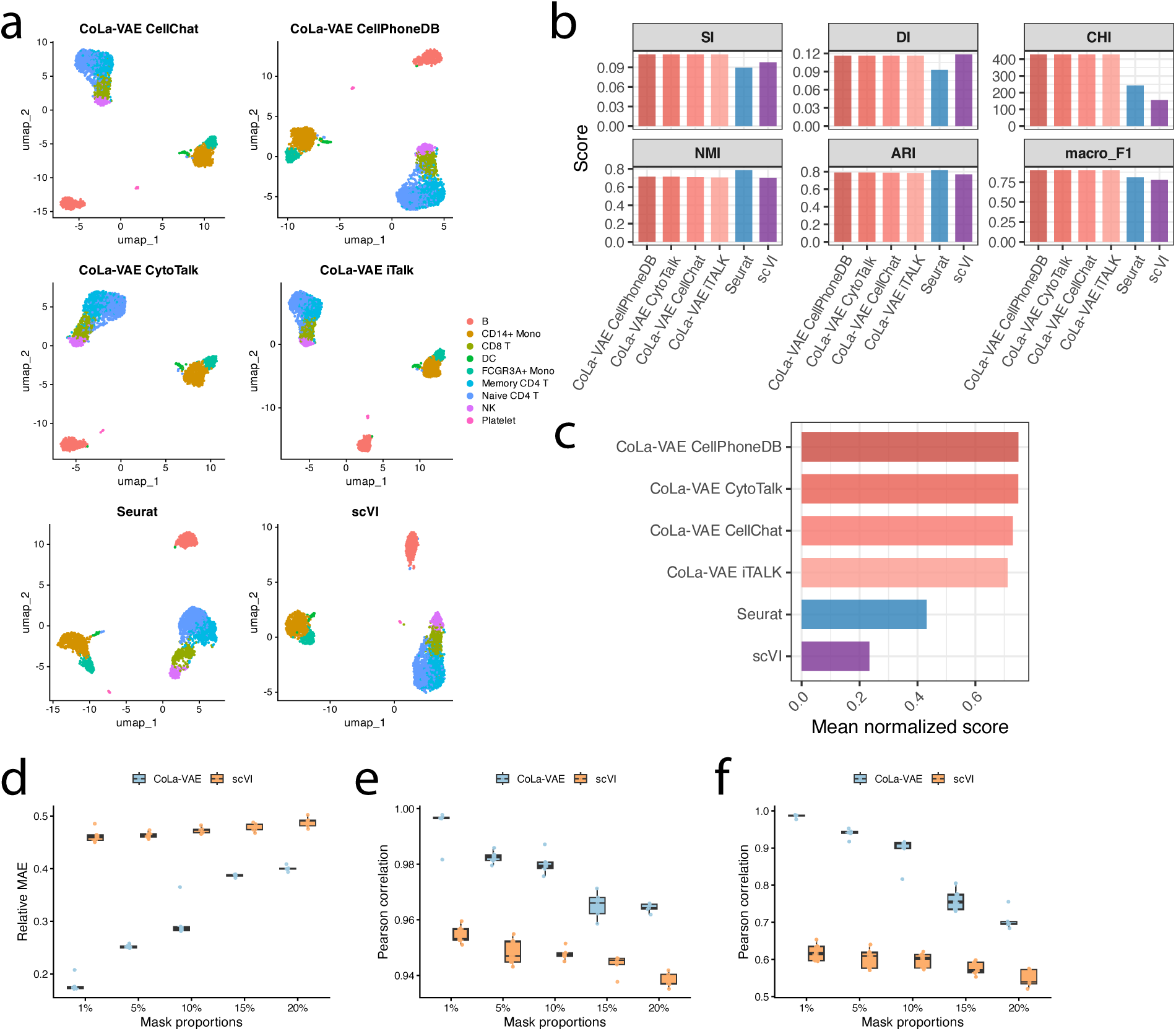
CoLa-VAE improves cell representation learning and robustly recovers masked gene expression in pbmc3k dataset. (a) UMAP visualization of latent embeddings generated by CoLa-VAE using CCC priors derived from different CCC inference methods (CellChat, CellPhoneDB, iTALK, and CytoTalk), compared with baseline methods Seurat and scVI. (b) Quantitative benchmarking of clustering performance across methods using six evaluation metrics, including silhouette index (SI), Davies-Bouldin index (DI), Calinski-Harabasz index (CHI), normalized mutual information (NMI), adjusted Rand index (ARI), and macro-F1 score. (c) Mean normalized performance scores summarizing the overall benchmarking results across all evaluation metrics. Result indicates CoLa-VAE achieved improved separation and organization of major celltypes. (d-e) Robustness evaluation of CoLa-VAE under different masking proportions (1%, 5%, 10%, 15%, and 20%) using five independent random seeds per condition. scVI as the baseline denoising method being compared. (d) Relative Mean Absolute Error (MAE) between recovered values and denoised values generated from the complete dataset. (e) Pearson correlation between recovered values and denoised values generated from the complete dataset. (f) Pearson correlation between CCC networks reconstructed from recovered data and those generated from the complete denoised dataset. These results demonstrating that CoLa-VAE can accurately recover masked gene expression and preserving global CCC structure.

To assess the contribution of the communication-aware latent component, we further compared the full CoLa-VAE model with latent subspaces and ablated variants. The CCC-specific latent dimensions alone retained substantial cell-type organization, whereas the normal expression-related dimensions and the model without CCC showed weaker separation for several cell populations (Supplementary Fig. 2a-c). Quantitatively, the full CoLa-VAE model achieved the highest overall score, while CCC-only, normal-only, and no-CCC variants showed reduced or more uneven performance across the 6 evaluation metrics (Supplementary Fig. 2d,e). These findings support that CCC-derived information contributes additional structure beyond the expression-only latent representation.

We also examined whether CoLa-VAE performance was sensitive to hyperparameter choices. Notably, across a broad range of normal and CCC latent-dimension allocations, performance did not monotonically improve with larger latent sizes. Instead, the best overall performance was observed when the total latent size was approximately 10–20 features, with CCC-specific dimensions accounting for about 20–30% of the latent space (approximately a 2:8 to 3:7 CCC-to-normal ratio; Supplementary Fig. 3a-g). This suggests that CoLa-VAE benefits from a compact latent representation with a balanced contribution from communication-aware and expression-related features, rather than from simply increasing latent dimensionality. In addition, varying the number of highly variable genes or ligand-receptor pairs did not show a monotonic improvement in performance, indicating that CoLa-VAE is relatively robust to moderate changes in feature selection (Supplementary Fig. 3h,i).

We then tested the robustness of CoLa-VAE denoising under increasing masking noise. After randomly masking expression values at proportions ranging from 1% to 20%, with lower-expression genes assigned a higher masking probability to mimic dropout-like sparsity, we compared the reconstructed outputs from CoLa-VAE and scVI across 5 independent random seeds. CoLa-VAE produced lower relative mean absolute error (MAE) than scVI across all masking levels, with the strongest advantage observed under low to moderate masking proportions (Fig. 2d). CoLa-VAE also maintained higher Pearson correlation with the reference expression profile than scVI (Fig. 2e). In addition, the communication profiles reconstructed from masked data remained more similar to the full-data reference in CoLa-VAE than in scVI (Fig. 2f). These results suggest that CoLa-VAE better preserves both expression-level information and downstream communication structure under perturbation.

Finally, we used SPARSim^13^ to evaluate denoising accuracy against ground-truth expression. 5 PBMC reference datasets with different cell numbers, library sizes, and sequencing technologies were used to generate simulated ground-truth expression matrices and corresponding count matrices. Across 5 random seeds per reference dataset, CoLa-VAE achieved consistently high Pearson and Spearman correlations with the simulated ground truth, outperforming scVI and 2 state-of-art imputation methods, MAGIC and ALRA (Supplementary Fig. 4a,b). Error-based metrics showed CoLa-VAE achieved low RMSE and MAE across datasets (Supplementary Fig. 4c,d), with gene-wise comparisons indicating improved or competitive performance relative to scVI, MAGIC, and ALRA (Supplementary Fig. 4e-f). Examination of recovered expression at masked positive and negative positions further suggested that CoLa-VAE preserved a clearer distinction between true signal and background than methods that tended to over-impute expression values (Supplementary Fig. 4g). Together, these simulation results provide direct support that CoLa-VAE denoised outputs more closely approximate underlying true expression.

### CoLa-VAE improves latent representation quality and denoising consistency across different single-cell sequencing platforms

To evaluate generalization across diverse technical conditions, we benchmarked CoLa-VAE using PBMC datasets generated from multiple sequencing platforms and experimental batches. These datasets included substantial differences in capture efficiency, sequencing depth, and dropout profiles, providing a challenging setting for representation learning and denoising. Across 9 PBMC datasets, structural clustering metrics, including SI, DI, and CHI and annotation-based metrics such as NMI, ARI, and macro-F1 remained comparable to other competing methods^14–17^ (Fig. 3a). When normalized scores were aggregated across all datasets and metrics, CoLa-VAE achieved the highest overall performance score among the competing methods (Fig. 3b).

**Figure 3.**
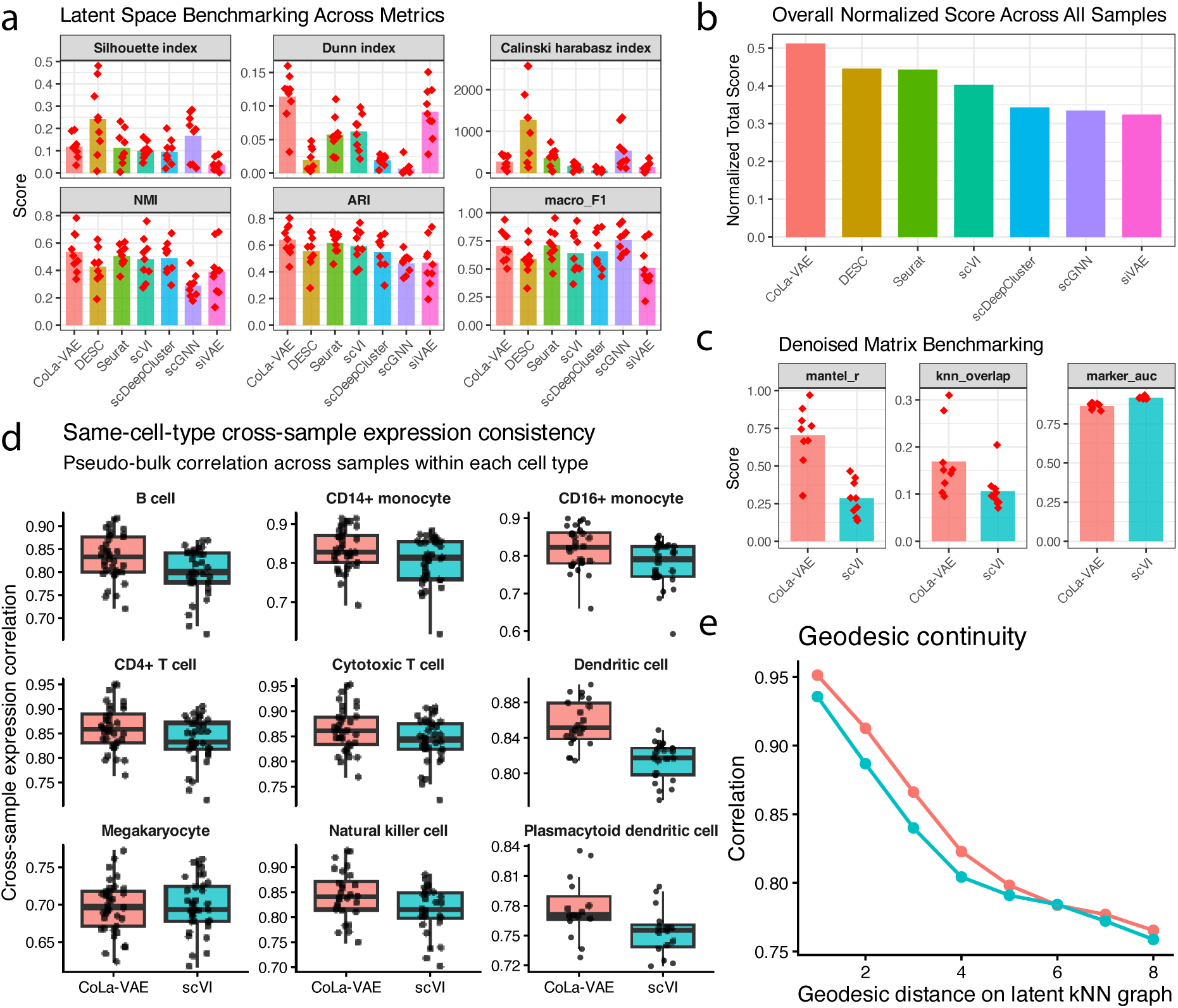
CoLa-VAE improves latent representation quality and denoising consistency across different single-cell sequencing platforms. (a) Benchmarking of latent space representations across 9 PBMC datasets generated from different sequencing platforms and experimental batches. Latent representations generated by CoLa-VAE and competing methods were evaluated using silhouette index (SI), Dunn index (DI), Calinski-Harabasz index (CHI), normalized mutual information (NMI), adjusted Rand index (ARI), and macro-F1 score. (b) Overall mean normalized performance scores aggregated across all benchmarking metrics and datasets. (c) Benchmarking of denoised expression matrices generated by CoLa-VAE and scVI using multiple evaluation metrics, including Mantel correlation, k-nearest-neighbor overlap, and marker gene AUC. (d) Cross-sample pseudo-bulk expression consistency analysis within each cell type. Spearman correlation coefficients were calculated between datasets generated from different sequencing platforms/batches for the same annotated cell type using denoised expression matrices from CoLa-VAE and scVI. (e) Geodesic continuity analysis measuring preservation of neighborhood relationships along the latent k-nearest-neighbor graph. CoLa-VAE maintained stronger local-to-global manifold continuity across increasing geodesic distances compared with scVI.

Because CoLa-VAE explicitly reconstructs a denoised expression matrix, we next assessed whether the denoised output preserved biologically meaningful structure. Compared with scVI, CoLa-VAE demonstrated improved Mantel correlation and k-nearest-neighbor overlap while maintaining comparable marker gene AUC values (Fig. 3c). These findings suggest that CoLa-VAE better preserves both global and local manifold structure during denoising without substantially compromising marker specificity.

To evaluate robustness across batches and sequencing protocols, we performed pseudo-bulk consistency analysis within each annotated cell type. Using denoised expression matrices, we computed cross-sample Spearman correlations between datasets generated from distinct experimental platforms. Across major immune cell populations, CoLa-VAE generally produced higher cross-sample concordance compared with scVI (Fig. 3d), supporting improved preservation of expression programs across different sequencing platforms. This increased cross-platform consistency is consistent with our simulation results showing that CoLa-VAE denoised matrices more closely approximate ground-truth expression, as the underlying biological expression program of a given cell type should be less dependent on sequencing technology than the observed noisy count matrix.

We additionally assessed geodesic continuity along the latent k-nearest-neighbor graph. CoLa-VAE maintained stronger correlation continuity across increasing geodesic distances compared with scVI (Fig. 3e and Supplementary Fig. 5), indicating improved preservation of local-to-global manifold organization after denoising.

Finally, to examine whether CCC-aware representations facilitate batch integration, we analyzed 3 independent PBMC datasets sequenced by 10x Chromium platform. Compared with Seurat and scVI, CoLa-VAE showed improved alignment of shared immune populations while retaining biologically meaningful structure, although residual batch-specific footprints remained detectable (Supplementary Fig. 6). These observations suggest that cell-cell communication is a kind of biological invariant and communication-derived constraints may provide biologically informative signals that are partially conserved across experimental batches.

### CoLa-VAE enhances robust detection of differential cell–cell communication in opioid use disorder dataset

We next investigated whether CoLa-VAE denoised matrices improve the sensitivity and reproducibility of differential CCC analysis in a human opioid use disorder (OUD) dataset^18^. Using single-nuclei transcriptomic profiles from control (n=22) and opioid-exposed individuals (n=22), we performed CCC inference using CellChat independently for each sample followed by population-level differential communication analysis (Fig. 4a).

**Figure 4.**
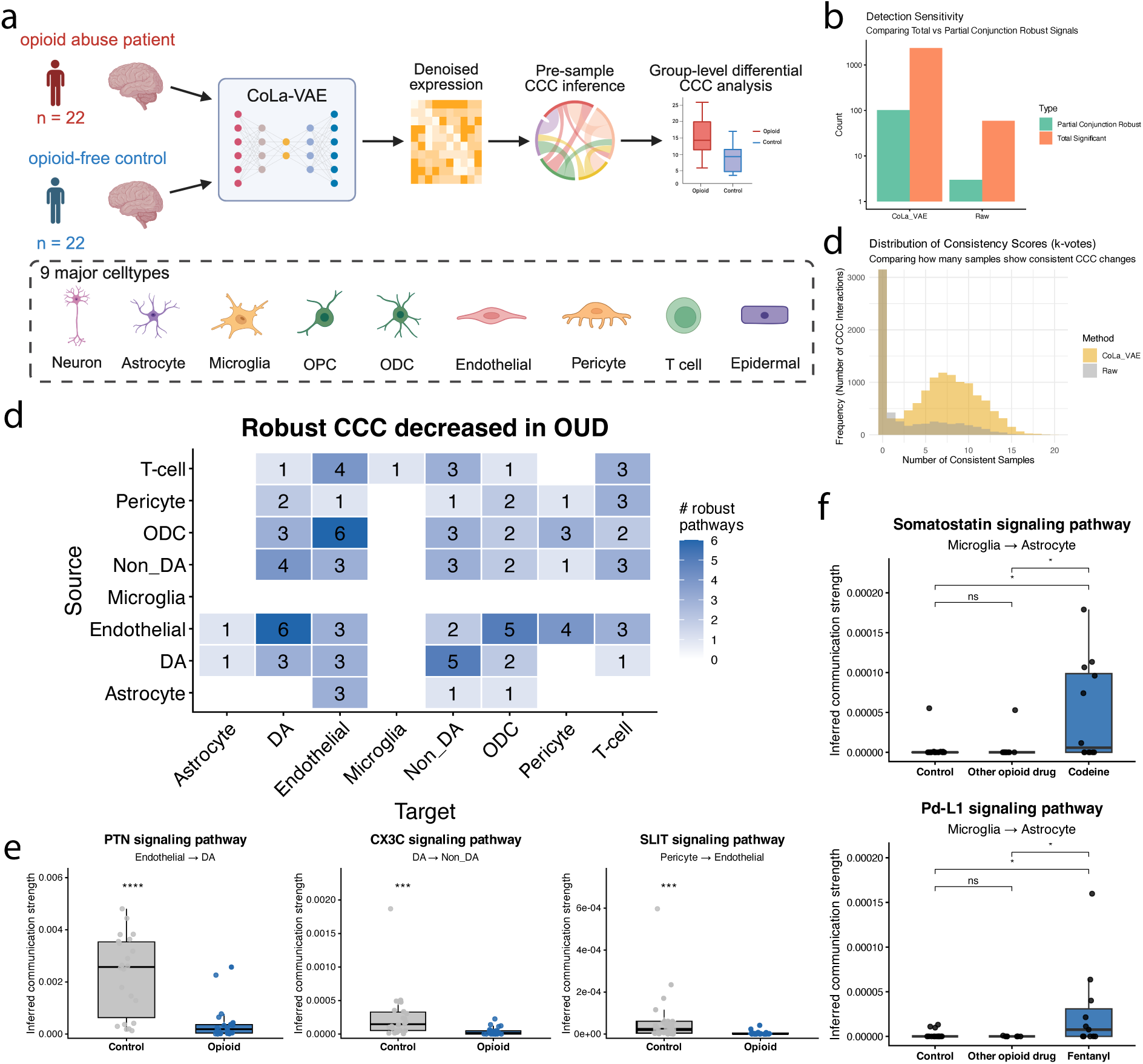
CoLa-VAE enhances the detection of robust and biologically consistent cell-cell communication alterations in opioid use disorder dataset. (a) Overview of the analytical framework for identifying differential cell-cell communication (CCC) in opioid use disorder dataset. (b) Comparison of differential CCC detection sensitivity between raw expression matrices and CoLa-VAE-denoised matrices. CoLa-VAE substantially increased the number of detected significant and partial-conjunction-robust CCC signals, indicating improved sensitivity and reproducibility of differential CCC analysis across samples. (c) Distribution of CCC consistency scores across samples, measured as the number of samples supporting concordant CCC alterations (k-votes). (d) Heatmap summarizing robustly decreased CCC pathways in OUD across source-target cell-type pairs. Numbers indicate the count of significantly decreased signaling pathways detected between each interacting cell pair. (e) Representative examples of significantly decreased signaling pathways in OUD, including PTN signaling from endothelial cells to dopaminergic neurons, CX3C signaling from dopaminergic to non-dopaminergic neurons, and SLIT signaling between pericytes and endothelial cells. Boxplots show inferred communication strength across control and opioid samples. (f) Drug-specific CCC alterations identified in opioid-exposed samples. Somatostatin and PD-L1 signaling from microglia to astrocytes showed selective increases associated with specific opioid compounds.

Compared with raw expression matrices, CoLa-VAE substantially increased the number of detectable significant CCC alterations as well as partial-conjunction-robust interactions supported across multiple samples (Fig. 4b, Supplementary Fig.7a). Consistency analysis further demonstrated that CoLa-VAE identified a larger number of recurrent CCC alterations shared across individuals (Fig. 4c), suggesting improved robustness against sparsity-related false negatives.

At the cell-type interaction level, multiple neuronal, glial, endothelial, and vascular communication programs exhibited coordinated decreases in OUD (Fig. 4d, Supplementary Fig.7b). Representative examples included reduced PTN signaling from endothelial cells to dopaminergic neurons, decreased CX3C signaling between dopaminergic and non-dopaminergic neuronal populations, and reduced SLIT signaling between pericytes and endothelial cells (Fig. 4e). These results indicate that OUD-associated transcriptional alterations are accompanied by widespread perturbations in intercellular signaling networks.

We additionally explored whether specific drug exposure was associated with distinct communication signatures. To identify candidate drug-associated CCC programs, we performed a LASSO regression screen in which inferred CCC programs were tested for associations with recorded drug exposures. Among the opioid-related signals, microglia-to-astrocyte somatostatin signaling was increased in codeine-positive samples, whereas microglia-to-astrocyte PD-L1 signaling was increased in fentanyl-positive samples (Fig. 4f). Although these associations are exploratory and require validation in larger cohorts, they suggest that CoLa-VAE may help identify drug-associated communication programs that are difficult to detect using sparse raw expression matrices alone.

### CoLa-VAE improves spatial transcriptomic deconvolution by incorporating spatial information

We next evaluated whether CoLa-VAE can be naturally extended to spatial transcriptomics, where the spatial proximity of cells provides direct biological context for cell–cell communication. Because CCC events are expected to occur preferentially between nearby cells, incorporating spatial information may help CoLa-VAE learn more accurate local communication profiles and generate denoised expression matrices that better preserve spatially resolved cellular composition.

To test this, we applied CoLa-VAE to a mouse brain spatial transcriptomics dataset^19^ with ground-truth cell type proportions provided by the original study. We compared MuSiC-based deconvolution results obtained from four input matrices: raw counts, CoLa-VAE denoised expression with spatial information, CoLa-VAE denoised expression without spatial information, and SpaVAE-denoised expression. The ground-truth dominant cell type map showed clear spatial organization of major neuronal and glial populations (Fig. 5a). When visualizing representative cell types, including Astro, Oligo, CA, Microglia, and L5_IT_CTX, deconvolution based on spatial CoLa-VAE more closely reproduced the ground-truth spatial probability patterns than raw counts or SpaVAE^20^ (Fig. 5b).

**Figure 5.**
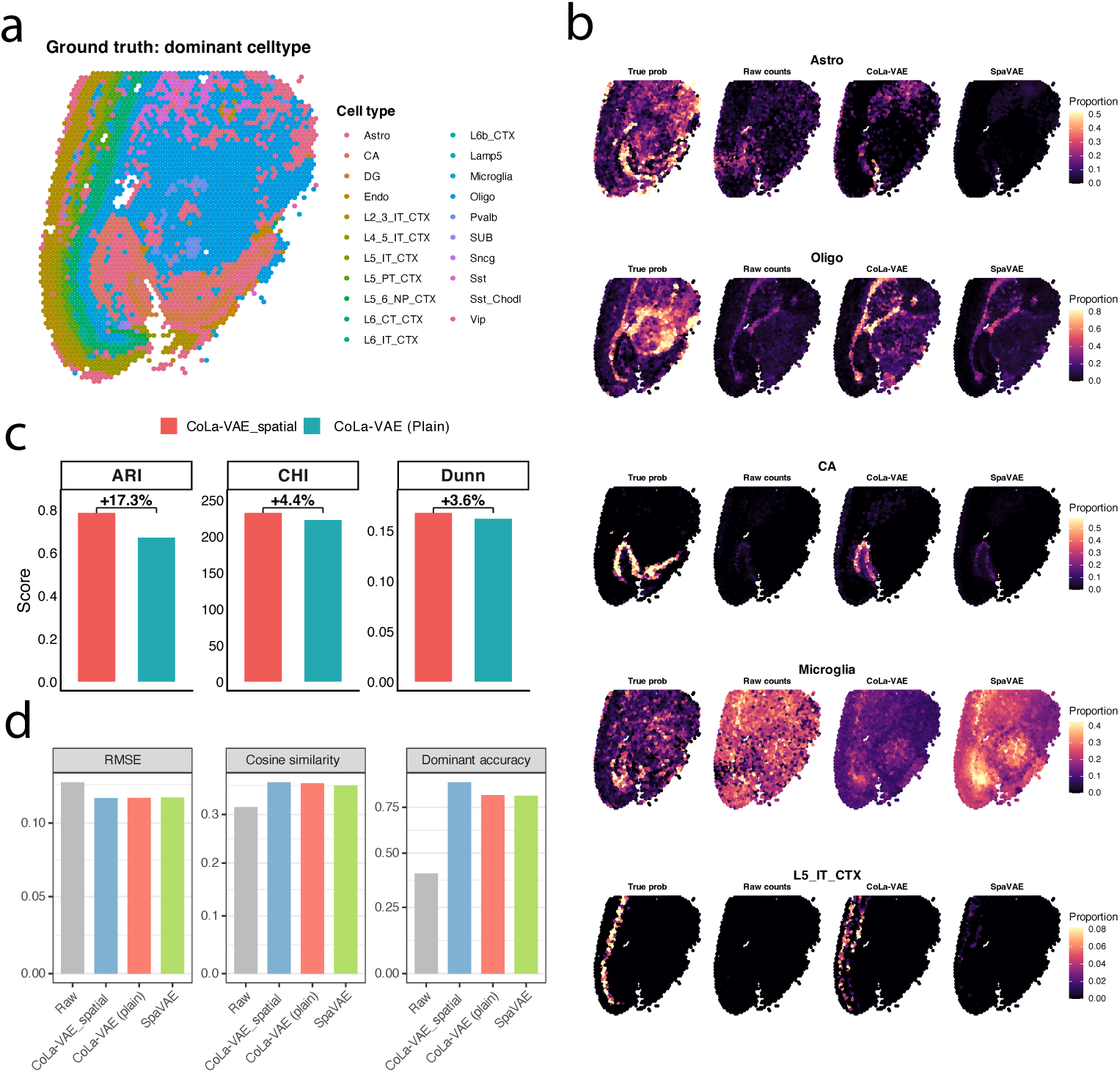
Spatially informed CoLa-VAE improves cell type deconvolution in mouse brain spatial transcriptomics. (a) Spatial distribution of the ground-truth dominant cell type for each spot in the mouse brain spatial transcriptomics dataset. Each spot is colored by the cell type with the highest ground-truth proportion. (b) Spatial maps of cell type proportions for five representative cell types: Astro, Oligo, CA, Microglia, and L5_IT_CTX. For each cell type, the ground-truth proportion is compared with MuSiC deconvolution results using raw counts, CoLa-VAE-denoised expression, and SpaVAE-denoised expression as input. (c) Comparison of latent clustering performance between spatial CoLa-VAE and plain CoLa-VAE without spatial information. Spatial CoLa-VAE improves ARI, CHI, and Dunn scores. (d) Quantitative evaluation of MuSiC deconvolution using raw counts, spatial CoLa-VAE-denoised expression, plain CoLa-VAE-denoised expression, and SpaVAE-denoised expression. Performance was assessed by RMSE, mean spot-wise cosine similarity, and dominant cell type accuracy.

Quantitatively, spatial CoLa-VAE achieved the best overall deconvolution performance across all 3 evaluation metrics, with the lowest RMSE, highest mean spot-wise cosine similarity, and highest accuracy in identifying the dominant cell type for each spot (Fig. 5d). In addition, incorporating spatial information improved the latent representation relative to the plain CoLa-VAE model, as reflected by higher ARI, CHI, and Dunn scores (Fig. 5c). Together, these results show that CoLa-VAE can be directly applied to spatial transcriptomics and that spatially informed CCC modeling improves both latent structure and downstream cell type deconvolution.

## Discussion

In this study, we introduced CoLa-VAE, a cell–cell communication-aware variational autoencoder that jointly learns latent representations and denoised expression profiles from sparse single-cell transcriptomic data. Although CoLa-VAE was originally motivated by the need to incorporate extrinsic signaling information into latent representation learning, our results indicate that one of its most important strengths is the quality and utility of its denoised expression output. Across masking experiments, simulation benchmarks, cross-platform PBMC datasets, disease-associated communication analysis, and spatial transcriptomic deconvolution, CoLa-VAE-denoised matrices consistently supported more robust downstream analysis than raw or competing denoised inputs.

A central observation from this work is that cell–cell communication provides a biologically meaningful prior for expression denoising. Conventional single-cell denoising and imputation methods primarily rely on expression similarity, local neighborhoods, or latent generative structure^21,22^. These strategies can be effective, but they may also smooth expression based on transcriptomic similarity alone, which can be confounded by technical noise, batch effects, or incomplete sampling of lowly expressed genes. CoLa-VAE introduces a complementary source of structure: cells are encouraged to share latent organization when they exhibit similar outgoing and incoming communication behaviors. Because ligand–receptor signaling reflects functional relationships among cells, this prior helps guide denoising toward expression patterns that are not only statistically plausible but also consistent with cellular interaction programs.

The simulation results provide direct support for this interpretation. Using SPARSim-generated datasets with paired noisy counts and ground-truth expression matrices, CoLa-VAE recovered expression profiles that were closer to the simulated ground truth than scVI and dedicated imputation methods such as MAGIC and ALRA. This result is important because it distinguishes denoising quality from downstream visual improvement alone. A denoised matrix can appear smoother or produce clearer clusters without necessarily being closer to the true expression state. By evaluating against simulated ground truth using correlation- and error-based metrics, we show that CoLa-VAE improves expression recovery rather than merely increasing apparent structure.

The downstream analyses further suggest that the denoised matrix is a central output of CoLa-VAE rather than a secondary byproduct. In the opioid use disorder dataset, CoLa-VAE-denoised expression increased the number of detectable and recurrent differential CCC signals compared with raw expression. This likely reflects improved recovery of ligand and receptor expression values that are missed or underestimated in sparse count matrices. Importantly, the additional signals were not interpreted solely as an increase in sensitivity; we also evaluated recurrence and partial-conjunction-supported interactions across samples, which provides evidence that the recovered CCC programs are more robust rather than simply more numerous. These results support the use of CoLa-VAE as a preprocessing and denoising step for downstream communication analysis, particularly in studies where sample-level or disease-associated CCC differences are of interest.

CoLa-VAE also improved denoised expression consistency across PBMC datasets generated from different sequencing platforms and experimental batches. Because the underlying biological expression program of a given cell type should be more conserved than the observed noisy count matrix, higher cross-platform pseudo-bulk concordance suggests that CoLa-VAE reduces some technology-dependent distortion while preserving cell-type-specific signal. We do not view CoLa-VAE as a replacement for dedicated batch integration methods, and residual batch-associated structure remained detectable in some analyses. However, these results suggest that communication-derived constraints may act as a partially conserved biological anchor, helping the model distinguish functionally meaningful expression programs from technical variation.

The spatial transcriptomic analysis further extends this principle. In spatial data, physical proximity provides direct context for potential cell–cell communication. By restricting communication structure according to spatial information, CoLa-VAE generated denoised expression matrices that improved downstream MuSiC-based deconvolution relative to raw counts, plain CoLa-VAE, and SpaVAE-denoised inputs. This result suggests that communication-aware denoising is particularly useful when the downstream task depends on recovering spatially organized cell type composition. More broadly, it supports the idea that denoising can be improved by incorporating biologically grounded constraints that reflect the structure of the tissue rather than relying on expression alone.

Several limitations remain. First, the quality of CoLa-VAE depends in part on the ligand–receptor database and communication scoring scheme used to construct the communication graph. Although we observed robust performance across several CCC formulations, incomplete or context-insensitive ligand–receptor annotations may limit performance in specific tissues or disease settings. Second, denoising and imputation can introduce artifacts if over-interpreted^21^. For this reason, CoLa-VAE-denoised matrices should be viewed as a complementary analytical layer rather than an unconditional replacement for raw counts, especially for differential expression analyses where count-level statistical assumptions are important. Third, our current spatial extension uses relatively simple spatial constraints. Future versions could incorporate more flexible spatial encoders or distance-aware graph neural networks to better model tissue architecture.

In future work, CoLa-VAE could be extended toward multi-sample and multi-modal integration by explicitly modeling communication programs that are conserved across individuals, technologies, or modalities. Such an extension would be particularly useful for large clinical cohorts, where the goal is not only to align cells across batches but also to preserve disease-relevant signaling programs and patient-specific biological variation. In addition, integrating CoLa-VAE with downstream uncertainty estimation may help distinguish confidently recovered expression values from ambiguous imputations.

Overall, our findings establish CoLa-VAE as a communication-guided denoising and representation learning framework for single-cell and spatial transcriptomic analysis. By using cell–cell communication structure to guide expression recovery, CoLa-VAE produces denoised matrices that more closely approximate underlying biological expression states and improve downstream analyses, including communication inference, cross-platform comparison, batch-effect mitigation, and spatial deconvolution. This dual role distinguishes CoLa-VAE from methods focused primarily on latent embedding quality and highlights the broader potential of biologically informed denoising for extracting reliable signals from sparse single-cell data.

## Methods

### The CoLa-VAE Framework Architecture

We developed CoLa-VAE, an intercellular communication-aware variational autoencoder that integrates cell-cell communication signals into latent representation learning through a graph Laplacian prior. By doing so, we ground the mathematical model in established biological principles of intercellular signaling, moving beyond purely statistical inference to capture the functional drivers of cellular identity.

### Latent Variable Model and Disentangled Latent Space

Let *X* = [*x*_1_, *x*_2_, …, *x_N_*] ∈ ℝ*^N^*^×*G*^denote the single-cell gene expression matrix, where *N* is the number of cells and *G* is the number of genes. Each cell *x_i_* is encoded into a latent embedding *z_i_* ∈ ℝ*^D^*. We decompose the latent variable as:

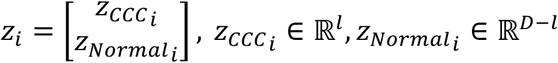

Here, *z_CCCi_* represents the CCC-aware latent dimensions that explicitly encode communication-driven topological structure, where *z_Normal_i__* captures residual transcriptional variation that associate with intrinsic cellular heterogeneity. We apply differential regularization to these subspaces: *z_CCC_* is subject to a graph Laplacian regularization prior derived explicitly from ligand-receptor interactions, while *z_Normal_* is subject to a Kullback-Leibler (KL) divergence constraint against a stand normal prior *N*(0, *I*). This disentangled design enables the model to separate biological signaling effects from generic gene expression heterogeneity.

The generative process of observed expression count *x_i_* in cell *i* is defined as:

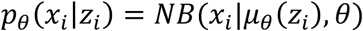

Where *μ_θ_*(⋅) is a neural decoder and *θ* denotes gene-specific dispersion parameters, The Negative Binomial likelihood naturally accounts for the over-dispersed count structure of scRNA-seq data and has been widely adopted in deep generative single-cell modes. The inference model *q*_ψ_(*z*|*x*) approximates the posterior distribution using a multivariate Gaussian parametrized by neural networks:

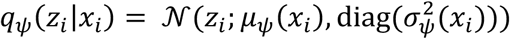

### Dynamic Inference of Cell-Cell Communication

Unlike methods that rely on static priors, CoLa-VAE employs a dynamic, iterative training strategy. At periodic intervals (every *T* epochs), we generate the denoised matrix *X^* (the decoder’s mean output) and use it to explicitly calculate pairwise cell-cell interaction scores. CoLa-VAE is modular, supporting four distinct scoring functions adapted from major CCC inference paradigms:

### Module CellChat: Mass Action Law

Based on the law of mass action, CellChat model the interaction probability for a ligand-receptor pair *k* between cell *i* and cell *j* using a Hill function formulation. This module explicitly accounts for the contribution of structural co-factor (agonists *AG* and antagonists *AN*):

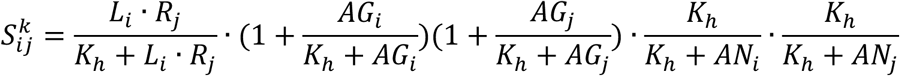

Where L, R, AG, AN are the geometric mean expression levels of the respective signaling molecules in the sender cell *i* or receiver cell *j*, if complex applicable. The *K*_ℎ_ is a saturation constant.

### Module CellPhoneDB: Multi-subunit Complexes

The model heteromeric protein complexes, CellPhoneDB define the functional expression of a complex as the minimum expression of its constituent subunits. For a ligand complex *L* and receptor complex *R*:

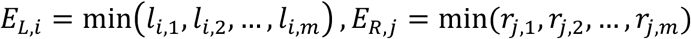

The interaction score is defined as the product of these function complex abundances:

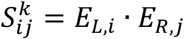

### Module iTalk: Product of Abundance

Following the iTalk methodology, we quantify signaling potential based on the log-transformed abundance of ligands and receptors. This formulation assumes that communication strength scales with the expression magnitude of the interacting partners:

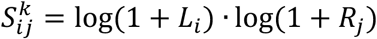

### Module CytoTalk: Mutual Information

This module prioritizes interactions with high statistical dependency. We first compute the Preferential Expression Measure (PEM) to define cell-type specificity. The interaction score combines the PEM with the Mutual Information (MI) between ligand and receptor expression distributions, prioritizing pairs that show non-random co-variation:

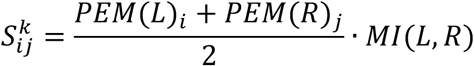

### Graph Construction via Bidirectional Communication Profiles

The raw pairwise interaction scores *S^k^_ij_* form an asymmetric matrix *P* ∈ ℝ*^N^*^×*N*^, where *P_i__j_* represents the signaling strength from cell *i* to cell *j*. To construct a symmetric adjacency graph suitable for Laplacian regularization, we measure the similarity between cells based on their global communication profiles rather than single edges.

We define the distance between two cells, *i* and *j*, by comparing their outgoing (sender cell) and incoming (receiver cell) signaling vectors. The Outgoing Distance *D_out_* measures the dissimilarity in how two cells signal to the rest of the population:

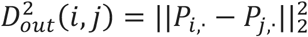

Similarly, the Incoming Distance *D_in_* measures the dissimilarity in how two cells receive signals from the population:

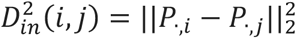

We combine these into a unifies Bidirectional Distance metric:

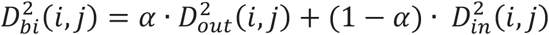

Where *α* (default 0.5) balances the contribution of sender and receiver profiles.

### Laplacian Regularization

Finally, we convert this distance into a Gaussian similarity kernel *W*:

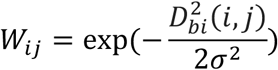

For spatial transcriptomics data, we additionally apply a spatial mask *M_spa_*, setting *W_i__j_* = 0 if the physical Euclidean distance between cells exceeds a radius threshold. The Normalized Graph Laplacian is then computed as:

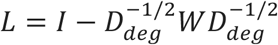

Where *D_deg_* is the degree matrix. This Laplacian L is used to compute the regularization loss ℒ*_Graph_*.

### ELBO Objective Function and Training Strategy

The final training objective is formulated as:

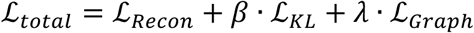

Reconstruction Loss (ℒ*_Recon_*) is the negative log-likelihood of the NB distribution, computed on a subset of Highly Variable Genes (HVGs):

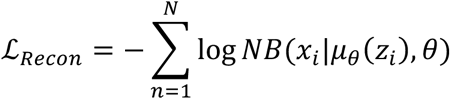

KL Divergence (ℒ*_KL_*) applied to *z_Normal_* to enforce the Gaussian prior:

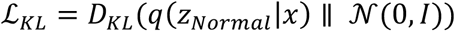

To prevent posterior collapse, we employed a Proportional-Integral-Derivative (PID) controller that dynamically adjusts *β* to maintain the KL divergency close to a target capacity.

Graph Laplacian Loss (ℒ*_Graph_*) applied to constrain the latent topology based on the dynamic communication graph:

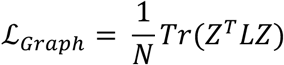

Here, *Tr*(⋅) denotes the matrix trace operator (sum of diagonal elements), computing the trace of this quadratic form effectively sums the Laplacian regularization cost across all *z_CCC_* latent dimensions. The model undergoes a warm-up period where graph regularization weight *λ* is set to zero initially. Subsequently, the communication graph L is recomputed every *T* epochs using the updated denoised expression *X^*, creating a positive feedback loop between reconstruction quality and topological inference.

### Benchmarking and Evaluation Metrics

To assess the quality of the learned embeddings, we performed Louvain clustering on the latent space representations and computed both structural and classification metrics. For structural metrics, we calculated the Silhouette Index (SI), Dunn Index (DI), and Calinski-Harabasz Index (CHI) to evaluate the compactness and separation of the resulting clusters without reference to ground truth labels. As for classification metrics, we compared the clustering results against ground truth annotations using Adjusted Rand Index (ARI) and Normalized Mutual Information (NMI). Macro-F1 Score is applied to robustly evaluate classification performance when the number of predicted clusters differs from the number of ground truth labels, we implemented a mapped Macro-F1 score. Each predicted cluster was assigned to the ground truth label that appeared most frequently within it (majority vote). We then calculated the F1 score for each class and reported the macro-average.

To evaluate the model’s ability to recover biological signals from noisy data, we benchmarked the CoLa-VAE-denoised matrices against baseline method (scVI) using three criteria. Global Structure Preservation (Mantel Test): We computed the Pearson correlation between the cell-cell distance matrices derived from the raw PCA space and the denoised PCA space. A higher Mantel statistic indicates better retention of global data geometry. Local Neighborhood Preservation (kNN Overlap): We identified the k=20 nearest neighbors for each cell in both raw and denoised representations. The overlap score was defined as the fraction of conserved neighbors, assessing the preservation of local manifolds. Biological Signal Retention (Marker AUC): We utilized canonical marker genes from PanglaoDB to quantify cell-type signal recovery. For each cell type, we calculated the mean expression of its markers in the denoised matrix and used this as a predictor score. We then computed the Area Under the Receiver Operating Characteristic Curve (AUC) for one-vs-rest classification of that cell type, averaging the AUCs across all annotated types.

### Expression-dependent masking strategy

Masking was performed on the count matrix of PBMC3k dataset. For each cell, only nonzero expression entries were considered, and cells with fewer than 10 nonzero genes were excluded from masking. Within each cell, masking probabilities were assigned as exp(-β\**x*), where *x* is the scaled expression value and the strength factor β was set as 2. This design gives lower-expressed genes a higher probability of being masked. For each masking ratio, a fixed fraction of nonzero entries in each eligible cell was sampled without replacement and set to zero. We generated masked datasets at five masking ratios: 1%, 5%, 10%, 15%, and 20%, with five independent repeats for each ratio. The original values and positions of the masked entries were recorded and used for downstream recovery evaluation.

### Simulation method

To benchmark denoising performance against simulated ground truth, we generated synthetic single-cell RNA-seq datasets using SPARSim. 5 10x Genomics PBMC datasets from TENxPBMCData were used as reference datasets, including PBMC3k, PBMC4k, PBMC6k, PBMC8k, and PBMC5k-CITEseq. Each dataset was converted to a Seurat object, followed by SCTransform normalization, PCA, nearest-neighbor graph construction, and Louvain clustering using 30 principal components and a clustering resolution of 0.5. The resulting Seurat clusters were used as condition labels for SPARSim parameter estimation. For each reference dataset, SPARSim parameters were estimated from the count matrix and cluster-defined cell groups, and five independent simulations were generated using different random seeds. The simulated outputs were saved and used for downstream denoising evaluation, with the simulated count matrix serving as the noisy observation and the corresponding simulated gene expression matrix serving as the reference ground truth.

## Data availability

All PBMC datasets can be downloaded via Seurat-data and TENxPBMCData R package. The Opioid dataset is available under accession number GSE240457.

## Code availability

The analysis and visualization scripts for this study are available at GitHub repository (https://github.com/Yeqing95/CoLa-VAE), along with a brief guide on reproducing the analysis workflows and figures presented in the paper.

## Acknowledgements

During the preparation of this work, the authors used Gemini and ChatGPT in order to improve readability and refine language. After using this tool, the authors reviewed and edited the content as needed and take full responsibility for the content of the publication. The authors thank X. Zhu from University of Pennsylvania for their feedback and discussions contributing to this work.

## Additional Information

Supplementary Information is available for this paper.

## Author contributions

Y.C. and Z.W. conceived and designed the study. Y.C. collected, and analyzed, and interpreted the data under the supervision of Z.W.. C.Q. and S.A. contributed to data interpretation and model design. F.L. and Z.Z. contributed to run competing method in benchmarking experiments. Y.C., C.Q. and H.F. contributed to manuscript preparation and revision. Y.C. wrote the manuscript with input from all the authors.

## Supplementary Figures

**Supplementary Figure 1.**
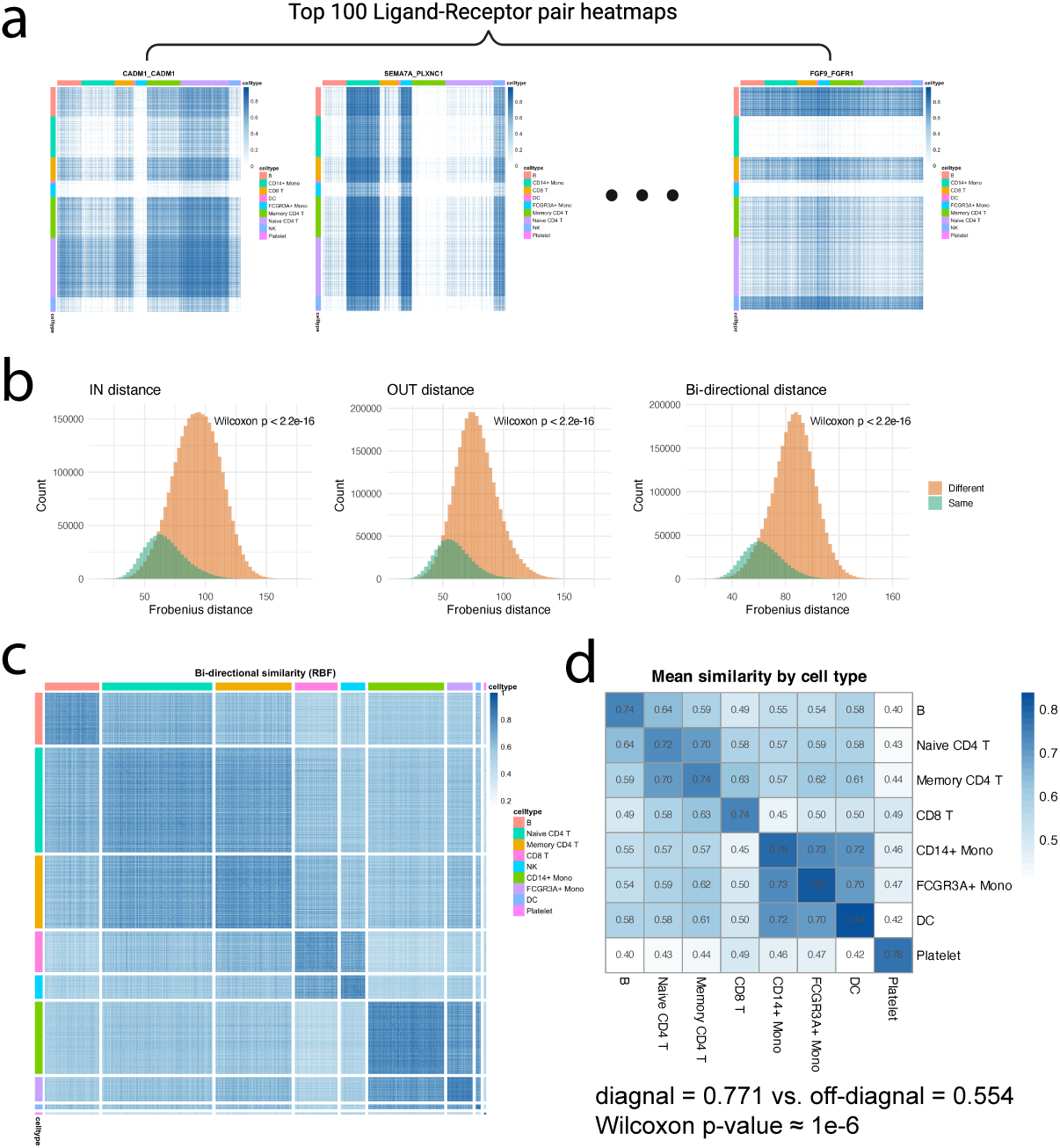
CCC-derived similarity structure in pbmc3k data. **(a)** Representative ligand–receptor (LR) pair communication matrices computed from denoised expression data. Each heatmap shows the strength of communication between pairs of cells for a given LR interaction. Block-like patterns indicate structured signaling relationships among groups of cells. **(b)** Distributions of pairwise Frobenius distances between cells based on outgoing, incoming, and bi-directional communication profiles. Cell pairs of the same annotated cell type (“Same”) exhibit significantly smaller distances compared to pairs from different cell types (“Different”), supporting that CCC profiles capture biologically meaningful similarity (Wilcoxon rank-sum test). **(c)** Cell–cell similarity matrix derived from aggregated LR communication signals using a radial basis function (RBF) transformation. Cells are ordered by annotated cell type. Enhanced block structure along the diagonal indicates that cells within the same type share similar communication patterns. **(d)** Mean similarity aggregated at the cell type level. Each entry represents the average similarity between two cell types. Higher values along the diagonal confirm that CCC-derived similarity is enriched within cell types, demonstrating consistency between communication structure and known biological annotations.

**Supplementary Figure 2.**
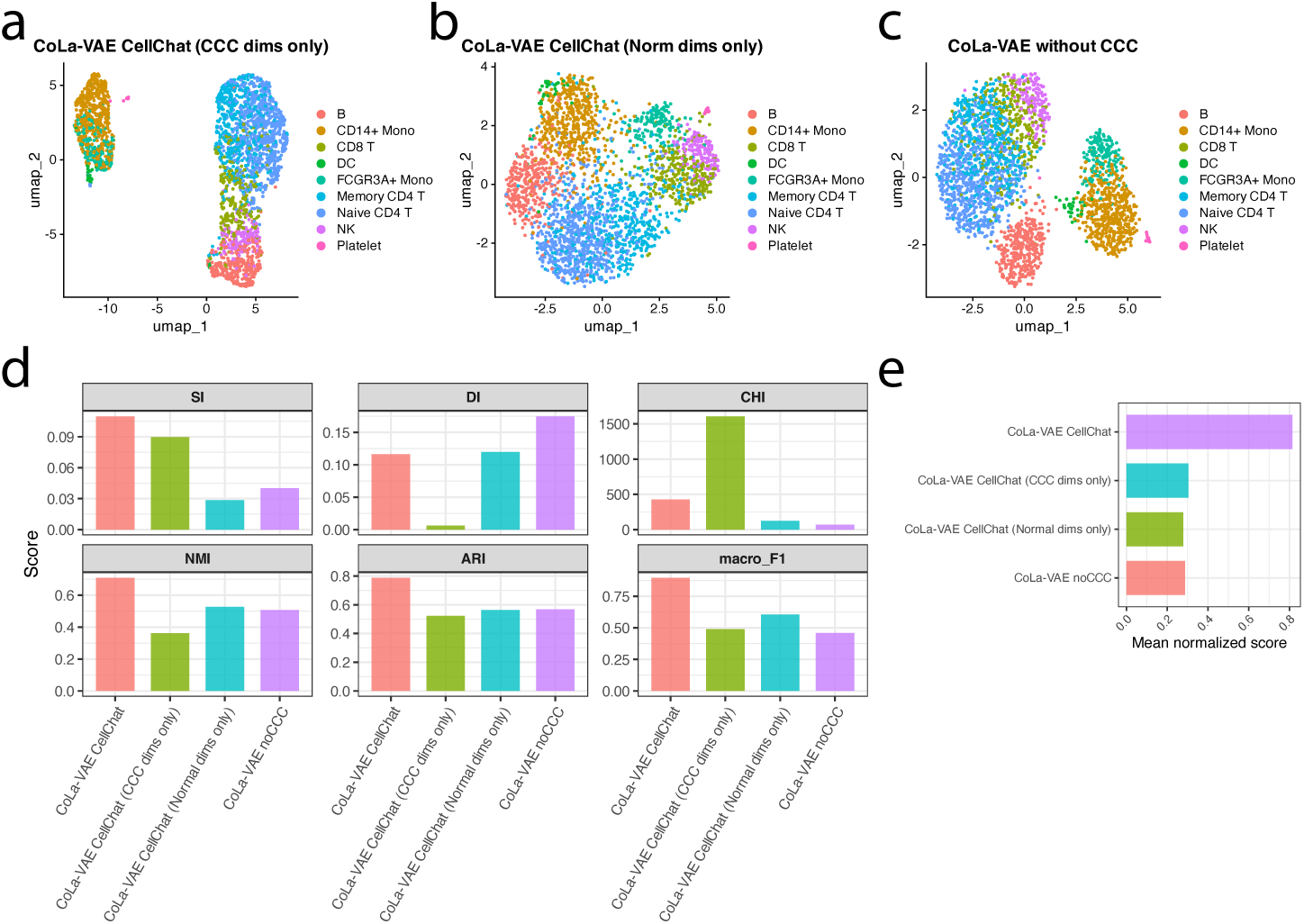
Comparison of CCC-specific and expression-specific latent representations learned by CoLa-VAE in pbmc3k dataset. (a-b) UMAP visualization generated using only the CCC-specific latent dimensions or only the normal expression-specific dimensions from CoLa-VAE trained with CellChat-derived communication priors. (c) UMAP visualization generated without using the CCC module from the same CoLa-VAE model. (d) Quantitative comparison of clustering performance using six evaluation metrics, including silhouette index (SI), Davies-Bouldin index (DI), Calinski-Harabasz index (CHI), normalized mutual information (NMI), adjusted Rand index (ARI), and macro-F1 score. (e) Mean normalized scores summarizing overall clustering performance of the full CoLa-VAE latent space, CCC-specific latent dimensions, and expression-specific latent dimensions. These results demonstrate that CCC-aware latent representations contribute substantially to improved cellular discrimination and biological structure preservation.

**Supplementary Figure 3.**
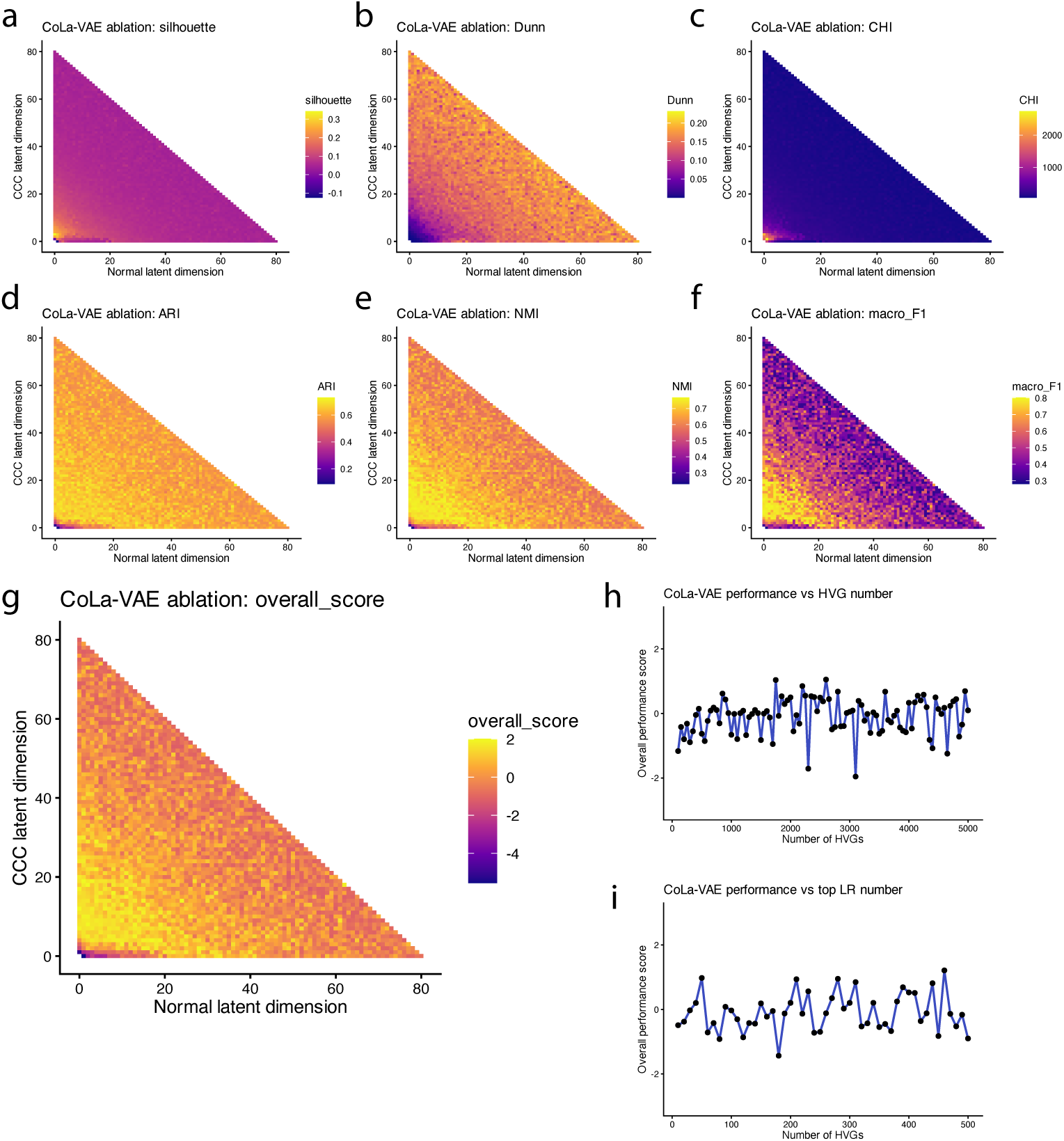
Ablation analysis of latent dimension allocation and input feature selection in CoLa-VAE in pbmc3k dataset. (a–g) Systematic ablation analysis evaluating the effect of different allocations between normal expression-related latent dimensions and CCC-specific latent dimensions on clustering performance. Heatmaps display performance across combinations of latent dimension sizes using multiple evaluation metrics, including silhouette score (a), Dunn index (b), Calinski-Harabasz index (CHI) (c), adjusted Rand index (ARI) (d), normalized mutual information (NMI) (e), macro-F1 score (f), and the aggregated overall performance score (g). Results demonstrate that balanced incorporation of CCC-specific latent dimensions consistently improves representation quality and clustering performance. (h-i) Sensitivity analysis of CoLa-VAE performance across different numbers of highly variable genes (HVGs) or varying the number of selected ligand-receptor pairs used for CCC modeling. Overall clustering performance remained stable across a broad range of HVG numbers or different CCC feature sizes, indicating robustness of CoLa-VAE modeling.

**Supplementary Figure 4.**
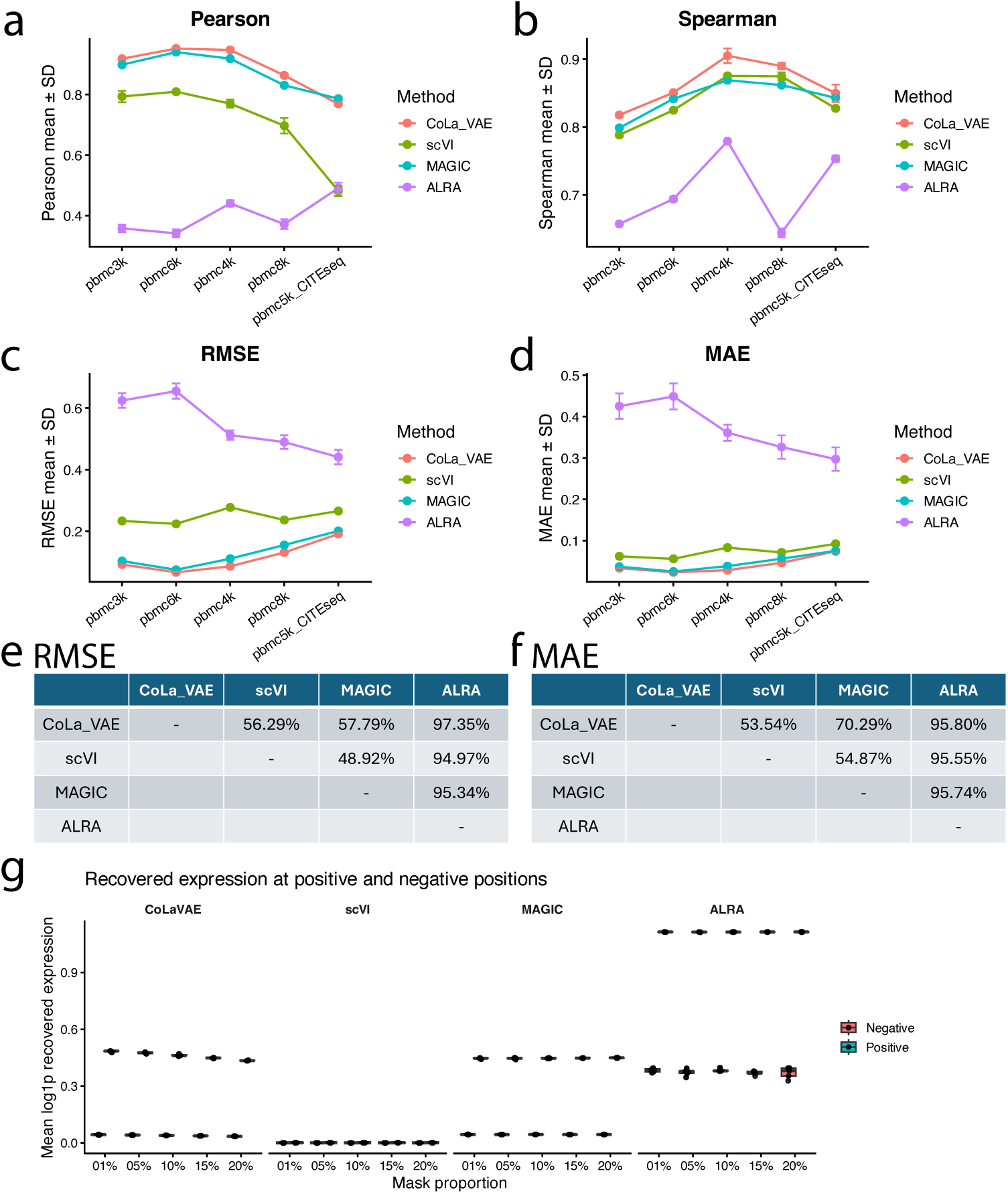
CoLa-VAE denoising accurately recovers ground-truth expression in simulated scRNAseq datasets. SPARSim was used to generate simulated scRNAseq datasets from five PBMC reference datasets with different cell numbers, library sizes, and sequencing technologies. For each reference dataset, 5 independent simulations were generated using different random seeds. The simulated count matrices were used as input for CoLa-VAE, scVI, MAGIC, and ALRA, and the resulting denoised or imputed matrices were compared with the simulated ground-truth expression matrices. (**a–d)** Global reconstruction performance was evaluated by calculating Pearson correlation (**a**), Spearman correlation (**b**), root mean squared error (RMSE; **c**) and mean absolute error (MAE; **d**) between the recovered and ground-truth expression values. Points and error bars represent the mean ± SD across five random seeds. (**e,f**), Gene-wise comparisons of RMSE (**e**) and MAE (**f**) across methods. Each entry indicates the percentage of genes for which the row method achieved lower reconstruction error than the column method. (**g**) Masking analysis performed on the simulated datasets generated from the pbmc3k reference. Nonzero entries were randomly masked at five masking proportions (1%, 5%, 10%, 15% and 20%) across 5 random seeds. The recovered expression values at artificially masked positions were compared with values at originally zero positions after denoising or imputation. CoLa-VAE selectively recovered masked positive entries while maintaining low expression at originally zero positions, indicating structured recovery rather than uniform inflation of zero values.

**Supplementary Figure 5.**
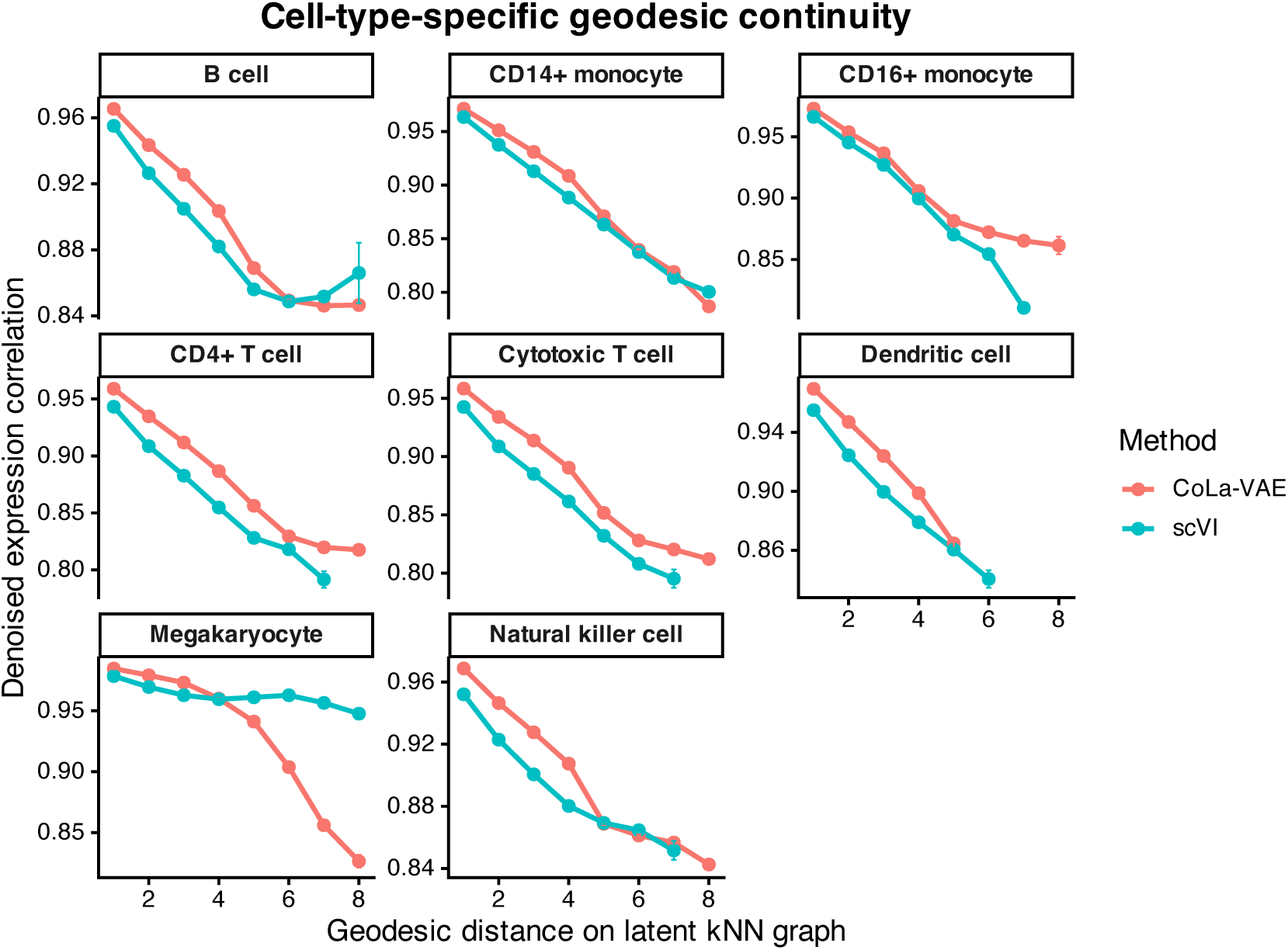
Cell-type-specific geodesic continuity analysis of denoised expression manifolds. Cell-type-specific evaluation of geodesic continuity in latent k-nearest-neighbor (kNN) graphs generated by CoLa-VAE and scVI across major PBMC cell populations. For each cell type, denoised expression correlation was measured as a function of geodesic distance along the latent manifold.

**Supplementary Figure 6.**
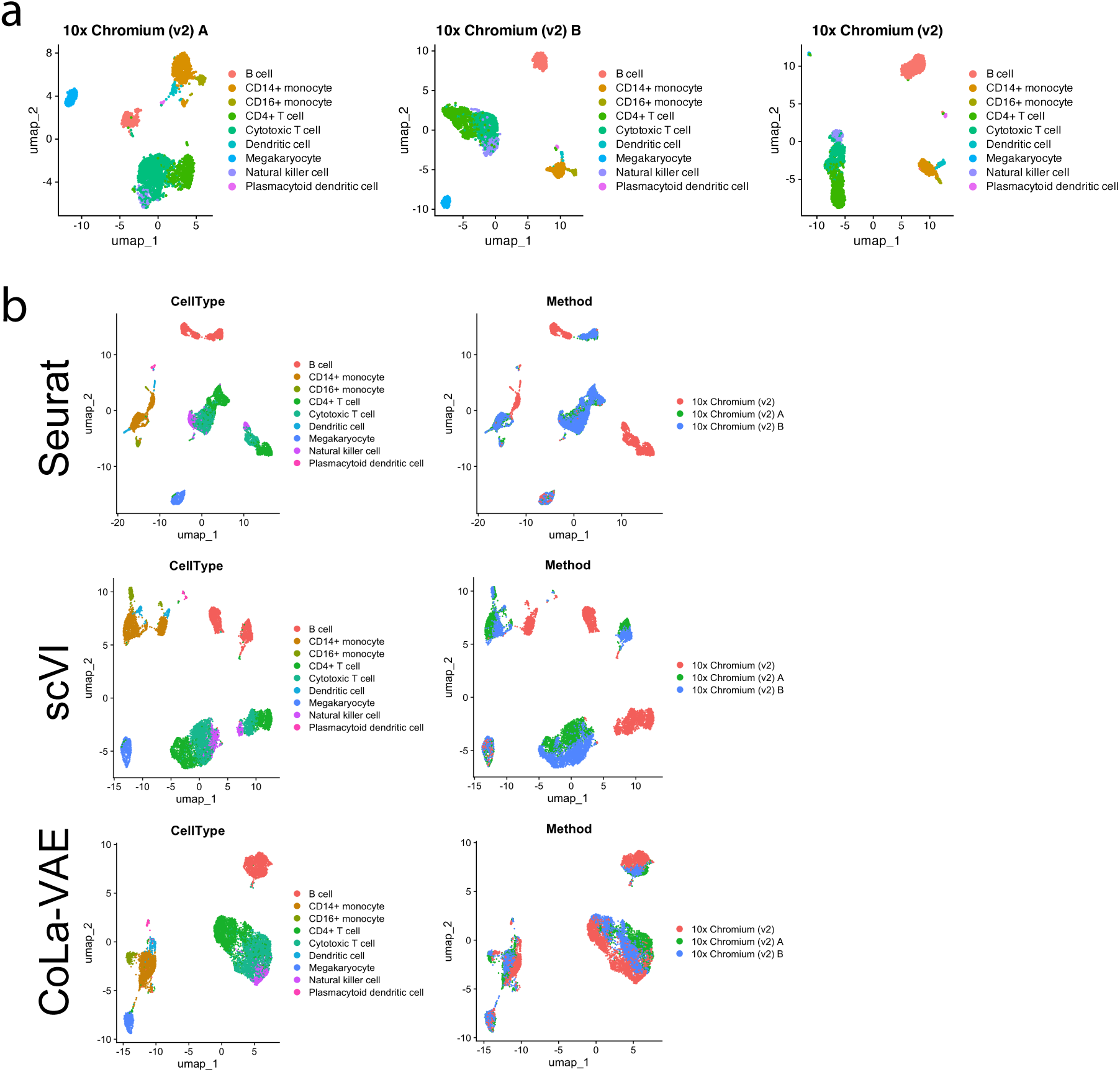
CCC-guided latent representations improve batch integration across independent PBMC datasets. (a) UMAP visualizations of three independent 10x Chromium PBMC datasets, showing consistent immune cell populations across datasets prior to integration. (b) Comparison of batch integration performance using Seurat, scVI, and CoLa-VAE. Left panels show embeddings colored by annotated cell type, while right panels show embeddings colored by dataset origin.

**Supplementary Figure 7.**
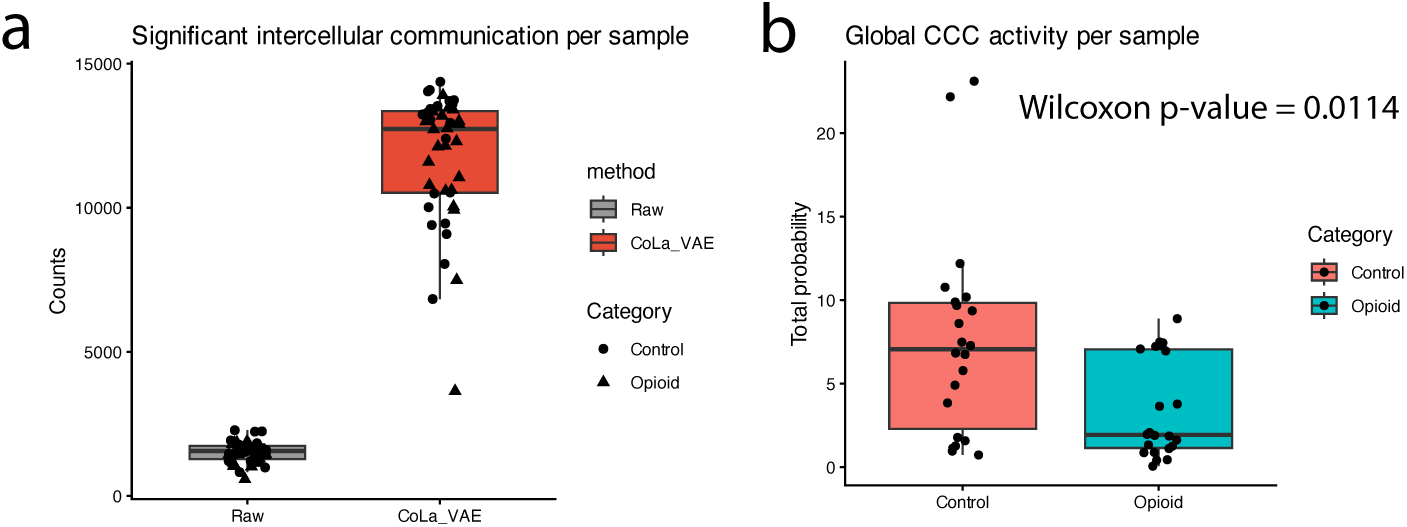
Global intercellular communication activity inferred from raw and CoLa-VAE-denoised expression matrices. (a) Total number of significant intercellular communication interactions detected per sample using raw and CoLa-VAE-denoised expression matrices. (b) Global CCC activity per sample measured as the summed communication probability across all inferred interactions.

